# Single-molecule turnover dynamics of actin and membrane coat proteins in clathrin-mediated endocytosis

**DOI:** 10.1101/617746

**Authors:** Michael M. Lacy, David Baddeley, Julien Berro

## Abstract

Actin is required for clathrin-mediated endocytosis (CME) in yeast. Experimental observations indicate that this actin assembly generates force to deform the membrane and overcome the cell’s high turgor pressure, but the precise molecular details remain unresolved. Based on previous models, we predicted that actin at endocytic sites continually polymerize and disassemble, turning over multiple times during an endocytic event. Here we applied single-molecule speckle tracking in live fission yeast to directly measure this predicted turnover within the CME assembly for the first time. In contrast with the overall ~20-sec lifetimes of actin and actin-associated proteins in endocytic patches, we detected single-molecule residence times around 1 to 2 sec, and high turnover rates of membrane-associated proteins in CME. Furthermore, we find heterogeneous behaviors in many proteins’ motions. These results indicate that rapid and continuous turnover is a key feature of the endocytic machinery and suggest revising quantitative models of force production.

## Introduction

Clathrin-mediated endocytosis (CME) is the major pathway for eukaryotic cells to internalize plasma membrane and extracellular molecules. In yeast, the invagination of a clathrin-coated pit (CCP) and formation of the ~50-nm vesicle relies on a dense meshwork of cytoskeletal actin filaments and associated proteins (Goode et al., 2015; Kaksonen et al., 2003; Kaksonen et al., 2006). This localized, rapidly-assembled actin meshwork is necessary to generate the force to bend the plasma membrane inward and overcome the cell’s high turgor pressure (Aghamohammadzadeh and Ayscough, 2009; Boulant et al., 2011; Collins et al., 2011). Although many endocytic proteins and their biochemical mechanisms have been characterized, our understanding of the molecular machinery is incomplete, especially in quantitative terms of how the endocytic assembly generates sufficient forces (Lacy et al., 2018).

CME is a challenging target for study because the entire assembly is smaller than the optical diffraction limit (~250 nm), the numbers of molecules at the site rapidly change, and most of the dynamics occur rapidly within ~20 sec. These unique challenges have made theoretical modeling and novel imaging techniques necessary for detailed study of CME (Berro and Lacy, 2018). Quantitative microscopy studies of the assembly and disassembly of endocytic proteins have revealed a robust timeline of self-assembly, as actin and associated proteins polymerize into a meshwork during invagination of the CCP and then gradually disassemble after the vesicle is pinched off (Berro and Pollard, 2014; Kaksonen et al., 2003; Picco et al., 2015; Sirotkin et al., 2010; Taylor et al., 2011). And while recent super-resolution imaging and electron microscopy of CME sites have revealed increasingly detailed structural organization (Arasada et al., 2018; Mund et al., 2018; Sochacki et al., 2017), these techniques are not currently able to quantify the dynamics of individual molecules within the endocytic site.

Theoretical calculations estimate that the amount of force needed to counteract turgor pressure and invaginate the membrane during CME in yeast is in the range of 1000-3000 pN (Carlsson and Bayly, 2014; Dmitrieff and Nedelec, 2015; Lacy et al., 2018; Tweten et al., 2017; Wang and Carlsson, 2017). The endocytic actin meshwork is composed of 100-200 short, Arp2/3-branched actin filaments that are capped shortly after nucleation; quantitative modeling indicates fewer than ten filament ends are polymerizing at any given time (Berro and Pollard, 2014; Sirotkin et al., 2010), each capable of producing around 1 pN of force (Footer et al., 2007; Kovar and Pollard, 2004). Even after accounting for other force-producing mechanisms, the force from a burst of actin polymerization appears to be insufficient to drive membrane deformation.

Based on the dendritic nucleation model and theoretical results (Berro et al., 2010), we hypothesized that continuous polymerization and disassembly of actin filaments underlies the observed dynamics in CME, which could allow the meshwork to generate more force than a single burst of filament polymerization. Turnover dynamics have been observed in other actin assemblies like lamellipodia (Pollard and Borisy, 2003; Watanabe and Mitchison, 2002; Yamashiro et al., 2014) and Fluorescence Recovery After Photobleaching (FRAP) experiments suggest the endocytic actin meshwork is highly dynamic (Kaksonen et al., 2003; Kaksonen et al., 2005; Picco et al., 2015). Continuous turnover of the endocytic actin meshwork components would allow the system to convert a larger amount of energy from ATP hydrolysis of actin polymerization into mechanical work over the meshwork’s lifetime. Turnover of network components like crosslinkers would also enable remodeling of the meshwork, allowing the redistribution of stored elastic energy (Ma and Berro, 2018; Picco et al., 2018) and higher order visco-elastic mechanisms. However, exchange and turnover of single molecules has not been directly observed in CME because conventional microscopy tools lack the resolution to measure such behavior within a transient and diffraction-limited structure like the endocytic patch.

In this study, we applied a variation of single-molecule speckle microscopy to track individual molecules in CME sites in live fission yeast. For eight target proteins, including actin-meshwork and membrane-coat proteins, we sparsely labeled the proteins in the cell and applied single-molecule localization and tracking analyses. These results provide the first direct evidence for rapid and continuous turnover of molecules in the endocytic actin meshwork on the timescale of 1-2 sec, and analysis of their motions reveals heterogeneous behaviors at the molecular level. Based on these results, we suggest that the amounts of force produced through known mechanisms might be higher than has been previously estimated.

## Results

### Different hypothetical mechanisms can generate similar bulk measurements

Previous studies have measured the numbers of molecules at endocytic sites, revealing characteristic robust profiles of assembly and disassembly over time (Berro and Pollard, 2014; Picco et al., 2015; Sirotkin et al., 2010). A common interpretation of those data is that the actin monomers and associated proteins polymerize into a meshwork during a discrete assembly phase accompanying the CCP invagination and then gradually disassemble after the vesicle is pinched off. However, studies of other actin systems have shown that filaments can continuously assemble and disassemble in a “treadmill” fashion (Roland et al., 2008; Watanabe and Mitchison, 2002; Yamashiro et al., 2014), with sustained turnover independent of the total number of molecules present. Based on the shared set of molecular components and theoretical models for endocytosis and other actin systems (Berro et al., 2010), we predicted that turnover might also underlie the apparent time-course of endocytic actin by altering the balance between nucleation/polymerization and disassembly to achieve net growth and net disassembly at different times.

We first performed simulations to demonstrate that such fundamentally different models as distinct assembly or continuous turnover could each generate apparent dynamics that resemble the previously observed microscopy data (Figure 1). In the simplest case, molecules are recruited and then disassembled in distinct phases (Figure 1A), resulting in the characteristic profile of growth and decay of the number of molecules at the site (Figure 1C, top row). A more specific simulation assumes that because polymerization of actin filaments predominantly occurs at their barbed end and depolymerization at the pointed end, the first molecules to be removed will be the oldest molecules (“first-in/first-out”, Figure 1C, middle row), which generates an identical profile of the number of molecules over time.

**Figure 1:**
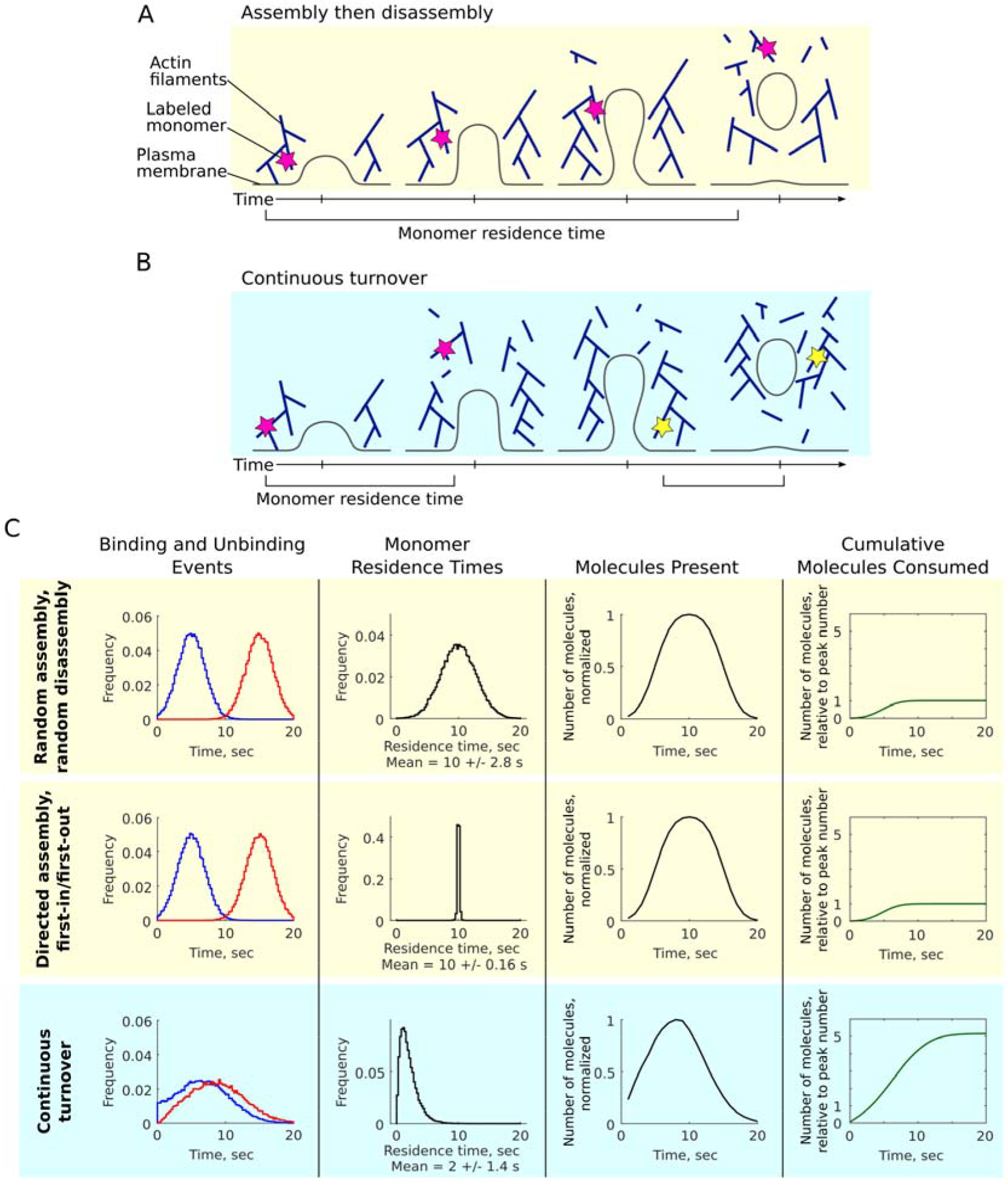
Different molecular mechanisms may give rise to similar apparent bulk dynamics. **A-B:** Many models suggest directed assembly and disassembly of actin in endocytosis but it is unclear whether the actin meshwork assembles and then disassembles in distinct phases (A) or continuously turns over during its bulk lifetime (B). Individual monomers are illustrated by magenta and yellow stars, with their residence times indicated. **C:** Simulated assembly and disassembly according to different hypothetical models: random assembly and disassembly in separate phases (top row), directed assembly with first-in/first-out disassembly (middle row), assembly with continuous turnover (bottom row). Far left: simulated occurrences of binding (blue) and unbinding events (red) underlying the assembly and disassembly mechanisms. Mid left: residence times of individual molecules, a quantity that clearly differentiates between the models. Mid right: number of molecules over time, a measurable quantity that can appear similar across the models. Far right: cumulative number of molecules consumed over time, normalized to the peak number of molecules present.

A model of assembly with continuous turnover of short-lived monomers (Figure 1B) can generate a profile of molecules over time that appears similar to the distinct assembly/disassembly models (Figure 1C, bottom row). The underlying differences in the models are illustrated by the binding and unbinding events (Figure 1C, left), features that are not directly accessible by experimental measurements. However, we might clearly distinguish these models by examining the residence times of individual molecules (Figure 1C, mid right). With random assembly and disassembly the residence times fall in a broad distribution with an average of half the total patch lifetime (10 sec), while for directed assembly with first-in/first-out disassembly, all molecules have identical residence times equal to half the patch lifetime (10 sec, with some noise). In contrast, continuous turnover results in a distribution of short residence times (2 sec). Although we have described these models in terms of actin monomers and filaments, similar behaviors could occur for other molecules recruited to oligomeric assemblies of any shape and organization.

Importantly, while the random assembly and first-in/first-out models consume only the number of molecules equal to the peak number of molecules present, the model of continuous turnover consumes a total number of molecules several times greater than the peak number (Figure 1C, right). The factor of total molecules consumed to peak number is simply dependent on the ratio of single-monomer residence time to patch lifetime (data not shown). To distinguish between these possible models, we sought to apply a single-molecule imaging strategy to directly measure the distributions of residence times in CME.

### Sparse labeling enables single-molecule imaging in endocytic sites

To measure single-molecule residence times in the dense assemblies of CME in live fission yeast cells, we employed a strategy based on single-molecule speckle microscopy (Watanabe and Mitchison, 2002; Yamashiro et al., 2014), an approach that we previously used to reveal dynamics of the yeast eisosome protein Pil1p (Lacy et al., 2017). We incubated cells expressing fusion proteins of the self-labeling SNAP-tag in media with low concentrations of a silicon rhodamine-647-conjugated SNAP substrate (SNAP-SiR) and imaged the cells in partial-TIRF. As reported in (Lacy et al., 2017), incubating live fission yeast with low concentrations of SNAP-substrate dye achieves only a low labeling efficiency (see Materials and Methods), enabling single-molecule tracking because typically one or zero of these sparsely labeled molecules are incorporated into diffraction-limited CME sites.

As a control to demonstrate that we are able to detect the full lifetime of both developing CCPs and vesicles diffusing after membrane scission in the partial-TIRF illumination field, we also tracked patches of fluorescently-tagged actin capping protein subunit (Acp1p-mEGFP, Figure 2A). We observed patches with an average lifetime of 18.4 +/− 3.5 sec (mean +/− S.D., Figure 2C, E, and Table 3), consistent with previous reports of capping protein lifetimes (Berro and Pollard, 2014; Sirotkin et al., 2010). We confirmed that the spots observed after SNAP-SiR labeling colocalize specifically with endocytic sites by imaging a strain co-expressing Acp2p-mEGFP and the endocytic actin crosslinker Fim1p-SNAP labeled with SNAP-SiR, and we observed negligible non-specific binding of SNAP-SiR dye in wild-type cells expressing no SNAP-tag (Supplemental Figure S1). In contrast to the many endocytic patches visible in Acp1p-mEGFP cells (70 to 120 endocytic events occurring per cell at any time (Berro and Pollard, 2014)), we observe only a few spots in cells expressing Acp1p-SNAP sparsely labeled with SNAP-SiR (referred to as Acp1p-SiR) (Figure 2B), typically 5 to 10 labeled molecules per cell per minute (Supplemental Figure S4E).

**Figure 2:**
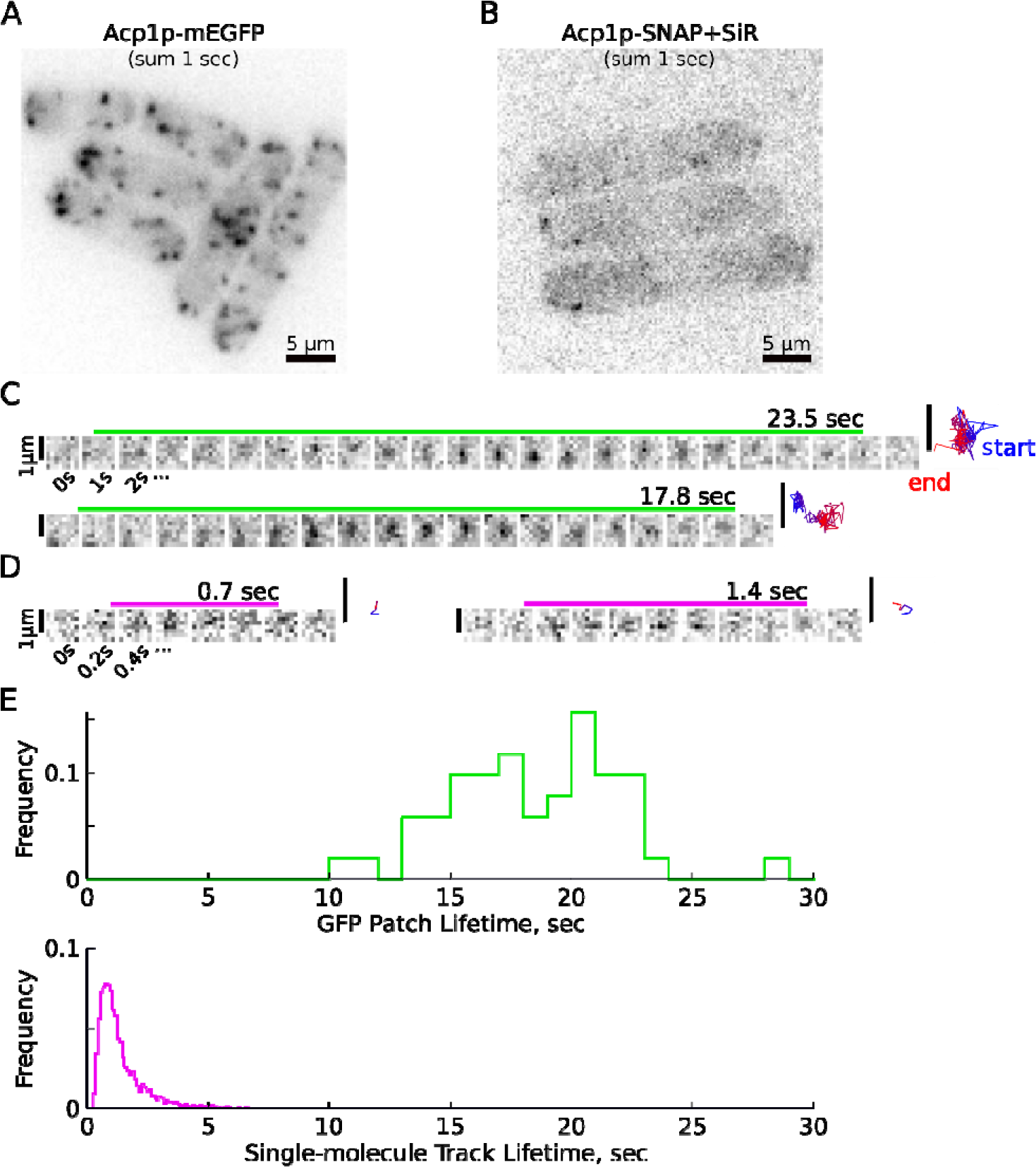
Sparse SNAP-tag labeling enables single-molecule speckle tracking of endocytic proteins. **A-B:** Sum projection images of 10 frames from a movie of Acp1p-mEGFP (A), or sparsely labeled Acp1p-SiR (B), inverted contrast. In A and B, scale bar is 5 µm. **C:** Montage of GFP spots, shown at 1 sec increments. **D:** Montage of SNAP+SiR spots, shown at 0.2 sec increments. In C and D, the lifetime of a spot is indicated by the colored bar above the image panels, and timescale is indicated below. At right is the trajectory of the spot or molecule, color coded with blue at the start and red at the end. Scale bars are 1 µm. Panels A-D have been prepared with different brightness settings for visual clarity. See supplemental Movies 1 and 2 for raw data. **E:** Top: Distribution of Acp1p-mEGFP patch lifetimes, as measured by semi-automatic tracking with TrackMate. N = 51 tracks from five movies recorded in two independent samples. Bottom: Distribution of Acp1p-SiR track lifetimes, as measured by single-molecule localization and tracking with PYME. N = 4,977 tracks from 24 movies recorded in five independent samples.

### Single-molecule residence times of endocytic proteins are short

We applied super-resolution localization and single-particle tracking algorithms (Baddeley et al., 2011) to the spots visible in SNAP-SiR labeled samples (see Supplemental Methods for further characterization of tracking results). Strikingly, Acp1p-SiR spots had short lifetimes in a peaked distribution with a long tail, with average 1.4 sec and 95% of events under 3.6 sec (Figure 2D-E). These single-molecule track lifetimes are much shorter than the 9 sec average predicted by the models of discrete assembly and disassembly (half of the Acp1p-mEGFP patch lifetime, as illustrated in Figure 1), instead consistent with the model of continuous turnover.

We imaged nine SNAP-tag strains and generated thousands of tracks for each target protein (Figure 3 and Table 1). The eight endocytic proteins included actin (Act1p), actin capping protein subunit (Acp1p), the actin crosslinker fimbrin (Fim1p), an Arp2/3 complex component (Arc5p), myosin-I (Myo1p), the nucleation-promoting factor Wiskott-Aldrich Syndrome Protein (WASp, Wsp1p), clathrin light chain (Clc1p), and the Hip1R/Sla2 homologue and actin-membrane anchoring protein (End4p). We also imaged a non-endocytic protein, the eisosome protein Pil1p, as a control protein that is expected to be immobile (Kabeche et al., 2011; Walther et al., 2006). Although a subset of Pil1p-SiR molecules undergoes rapid binding and unbinding at the ends of the eisosome structure (Lacy et al., 2017), a much greater fraction of Pil1p-SiR spots persists longer than 5 sec and the average lifetime is longer than any of the endocytic proteins (Figure 3 and Table 1).

**Table 1:** Statistics for single-molecule tracks of SNAP-tag fusion proteins.

| **Name** | **Lifetime**<br>**Median,**<br>**sec** | **Lifetime**<br>**Mean,**<br>**sec** | **Lifetime**<br>**S.D., sec <sup>(a)</sup>** | **Lifetime**<br>**95%ile,**<br>**sec** | **End-to-end**<br>**distance**<br>**Median, nm** | **N tracks** | **N movies**<br>**(datasets with**<br>**>40 tracks)** | **N**<br>**samples** | **D, nm<sup>2</sup>/sec <sup>(b)</sup>** |
| --- | --- | --- | --- | --- | --- | --- | --- | --- | --- |
| Act1p | 1.1 | 1.5 | 0.12 | 4 | 159 | 7,660 | 18 (18) | 2 | 2,955 |
| Acp1p | 1.1 | 1.4 | 0.19 | 3.6 | 167 | 4,977 | 24 (21) | 5 | 3,874 |
| Fim1p | 1 | 1.5 | 0.17 | 4.2 | 154 | 16,490 | 12 (12) | 2 | 2,855 |
| Arc5p | 1.2 | 1.6 | 0.32 | 4.2 | 166 | 4,576 | 17 (13) | 2 | 3,337 |
| Myo1p | 1.1 | 1.6 | 0.16 | 3.9 | 156 | 3,951 | 10 (10) | 2 | 1,678 |
| Wsp1p | 0.9 | 1.2 | 0.13 | 2.6 | 147 | 4,379 | 41 (32) | 5 | 1,869 |
| Clc1p | 1.1 | 1.5 | 0.16 | 3.7 | 168 | 2,718 | 19 (18) | 3 | 2,770 |
| End4p | 1.1 | 1.6 | 0.24 | 4.1 | 148 | 3,983 | 31 (17) | 5 | 1,381 |
| Pil1p | 1.2 | 2.0 | 0.67 | 5.9 | 134 | 446 | 6 (4) | 1 | 517 |

(a): Standard Deviation is determined across means calculated from individual datasets (each movie), only counting movies with > 40 tracks.
(b): Apparent diffusion coefficient calculated from mean squared-displacement over time for time windows < 1 sec (10 points), as in Figure 5.

**Figure 3:**
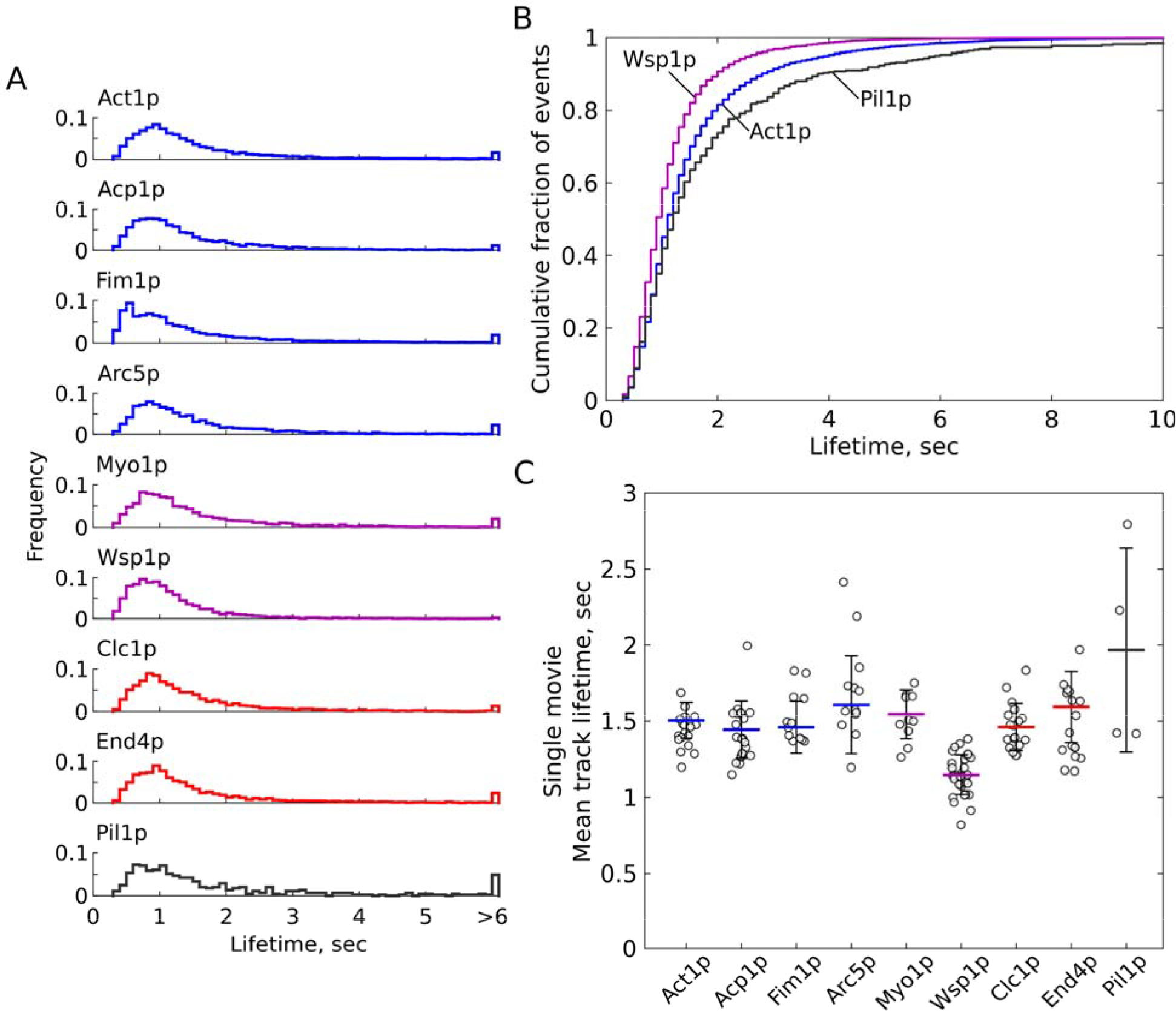
Single-molecule residence times of several endocytic proteins. Cells expressing SNAP-tag fusion proteins were sparsely labeled with SiR-647 then imaged and tracked as described in text. **A:** Probability distributions of track lifetimes for each target protein. Distributions are truncated at 6 sec and all events longer than 6 sec are shown in the final bin. The minimum allowed track length is 0.3 sec. **B:** Cumulative distributions of Act1p (blue), Wsp1p (purple) and Pil1p (gray). **C:** Mean track lifetimes calculated for each movie and standard deviation (calculated across means of all images with more than 40 tracks), individual dataset means shown by open circles. See Table 1 for summary statistics. Actin-associated endocytic proteins are colored in blue, nucleation-promoting factors are purple, and membrane-associated endocytic proteins are red. Pil1p is included as an “immobile” control in gray, although the detected tracks represent a mixture of stable molecules incorporated in eisosomes and dynamic molecules at eisosome ends.

Notably, single-molecule tracks of actin-associated proteins (Act1p, Acp1p, Fim1p, and Arc5p, which were all previously reported to have bulk lifetimes around 20-25 sec (Picco et al., 2015; Sirotkin et al., 2010)) display shorter lifetimes averaging 1.4 to 1.6 sec (Figure 3 and Table 1), while Wsp1p displays even shorter lifetimes (mean = 1.15 sec, 95%ile = 2.6 sec). End4p and Clc1p, which have been reported to have bulk lifetimes around 40 sec and 110 sec (Sirotkin et al., 2010) also display short average single-molecule residence times of 1.6 and 1.5 sec, respectively. All of the target proteins display a similar shape of residence time distribution, except for Fim1p, which contains a unique sub-population of short-lived events around 0.5 sec. Small but statistically significant differences occur between the distributions of lifetimes for many samples (by Mann-Whitney test, Supplemental Fig. S3), with Fim1p and Wsp1p significantly different from all other samples (at p < 0.0001). All samples except for Arc5p are significantly different from Pil1p (at p < 0.05). As with the comparison for Acp1p-mEGFP, all of the endocytic proteins we measured display much shorter lifetime than would be predicted for a simple assembly/disassembly model given their bulk lifetimes that have been reported previously (Berro and Pollard, 2014; Sirotkin et al., 2010).

We attribute the disappearance of spots to dissociation of proteins from endocytic sites, and not to photobleaching or other photophysical artifacts. While increasing illumination intensity caused a corresponding increase in the sample photobleaching rate as measured by the decay in the number of spots visible per frame (0.12 sec^−1^, about ten-fold slower than the track lifetimes we measure), the distributions of track lifetimes did not change significantly (Supplemental Figure S2). Additionally, in images of Acp1p-SiR cells fixed with paraformaldehyde, we could track many fluorescent spots over 20 sec long under the same illumination conditions (Supplemental Figure S2C). Therefore, while photobleaching does occur during the imaging time (~1 min), it is not occurring on the fast timescales (~1 sec) that would interfere with our ability to track single fluorophores.

### Characteristic motions of endocytic proteins are heterogeneous

The single-molecule tracks typically travel a net distance of only 100 to 200 nm (Figure 4A). Actin-associated proteins and clathrin contain a greater proportion of long tracks (> 250 nm, Figure 4B), while Pil1p tracks cover the shortest distances. We investigated the relationship between track lifetime and net distance traveled. For molecules undergoing pure diffusive motion, the end-to-end distance increases with the square root of the lifetime. However, we observed end-to-end distances depend on lifetime only for some proteins (Figure 4C). Comparing the tracks in the top and bottom 10% of lifetimes shows that long-lived tracks of actin-associated proteins (Act1p, Acp1p, Fim1p, and Arc5p) travel farther than short-lived tracks (Figure 4B). For membrane-bound endocytic proteins (Myo1p, Wsp1p, End4p, and to some extent, Clc1p) this dependence is lost (Figure 4B). Pil1p displays overall short displacements with a median track length of 134 nm, likely limited by the localization precision.

**Figure 4:**
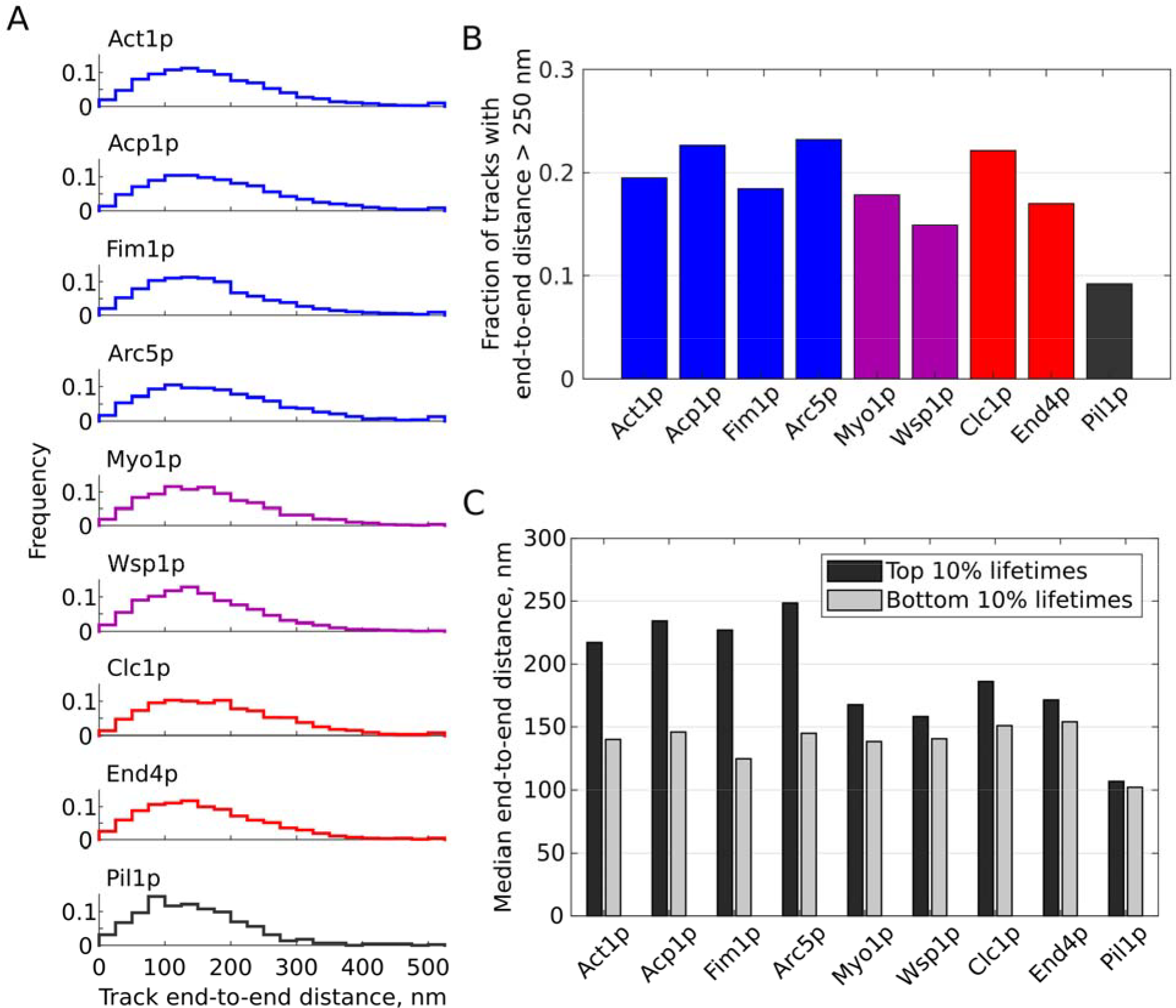
Single-molecule tracks net distances. **A:** Distributions of end-to-end distances for tracks of SNAP-tag fusion proteins. Actin-associated endocytic proteins are shown in blue, nucleation-promoting factors in purple, membrane-associated endocytic proteins in red, Pil1p as an “immobile” control in gray. **B:** Fraction of tracks with end-to-end distance longer than 250 nm for each sample. Colors as in (A). **C:** Median distances of tracks in the top or bottom 10% of lifetime distributions for each sample.

Next, we characterized each protein’s average mobility by calculating the mean squared displacement (MSD) over time (Figure 5). For all target proteins the MSD is linear at short time scales, suggesting diffusive behavior, but appears sub-diffusive at longer timescales (not shown here, but see Figure 6). We used only the first 10 points (up to 1 sec) for fitting, relying on only the highest-confidence points and avoiding potential biases resulting from the spot localization precision and lower number of points at long time steps (Michalet, 2010), and avoiding the regime of sub-diffusive motion. The slopes of these curves were significantly different (Figure 5A), but the y-intercept values are related to the localization precision (Michalet, 2010) which was similar across all samples (Supplemental Figure S3A). We computed the apparent diffusion coefficient D (Figure 5B), as 〈*dr*^2^〉 = 4*D* ⋅ *dt* + *b*, assuming a two-dimensional diffusion model because our images record only a single plane near the base of the cell, not three dimensions. Tracks of actin-associated proteins and clathrin have higher apparent mobility (2,700 to 3,800 nm^2^/sec) than those of membrane-associated proteins Myo1p, Wsp1p, and End4p (1,300 to 1,800 nm^2^/sec). However, these less-mobile endocytic proteins still display higher average mobility than Pil1p (~500 nm^2^/sec).

**Figure 5:**
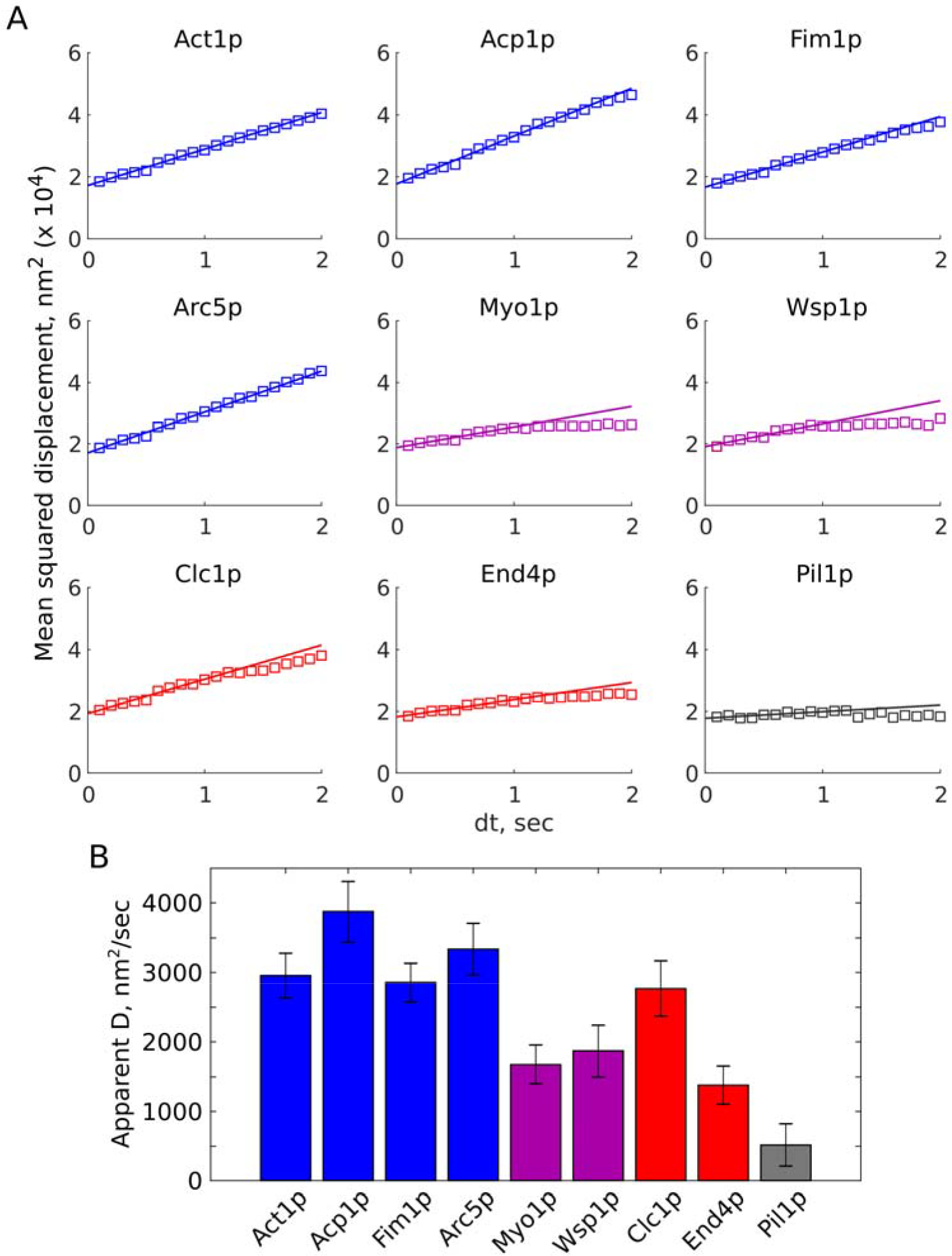
Average motions of endocytic proteins. **A:** Mean squared-displacement over time for all tracks (squares) with their linear fits (solid lines). **B:** Apparent diffusion coefficients for analyzed proteins, assuming 2-dimensional diffusion, MSD = 4D*dt + b. Error bars show the 95% confidence interval associated with the fit of MSD vs. dt slope. Actin-associated endocytic proteins are shown in blue, nucleation-promoting factors in purple, membrane-associated endocytic proteins in red, Pil1p as an “immobile” control in gray.

**Figure 6:**
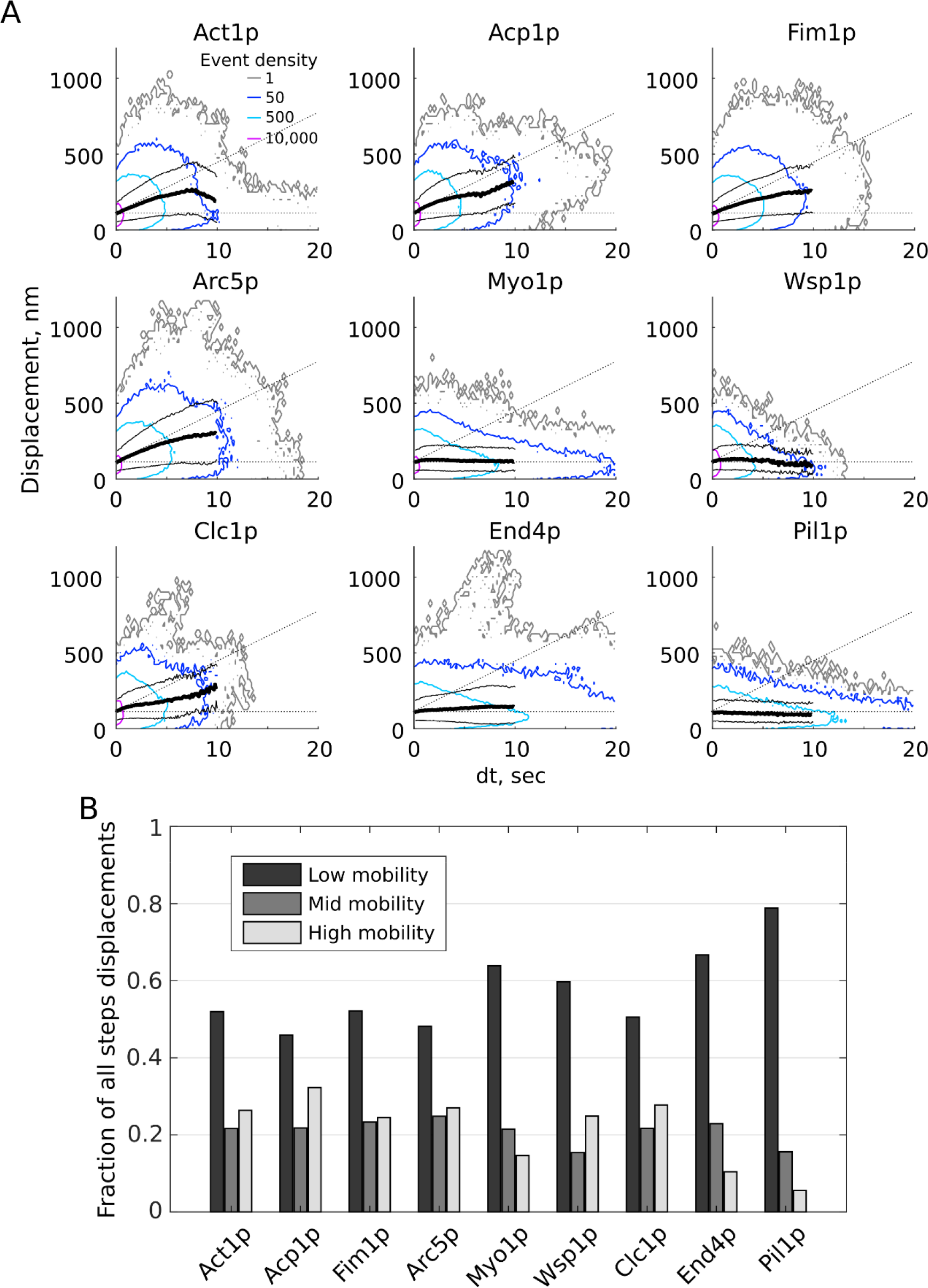
Distributions of single-molecule motions of endocytic proteins. **A:** All pairwise distances and time steps were calculated for tracks in each dataset, then binned and plotted in a 3D histogram. Contour lines represent bins of identical height, normalized by the total number of points in each dataset, with relative density (arbitrary scaling units) as indicated in the first panel. The mean displacement (heavy black line) and standard deviation (thin black lines) are shown for each time step below 10 sec. The high-, mid-, and low-mobility zones are indicated by black dotted lines. **B:** The fractions of all pairwise displacements from (A) that fall into high-, mid-, and low-mobility classes for each dataset.

We observed a high degree of heterogeneity in the full distributions of stepwise displacements (Figure 6A), and apparent differences in the shape of the distribution between proteins. To identify classes of behaviors, we divided each plot into areas of high, middle, and low mobility and determined the fraction of steps in each category for each protein (Figure 6B). The cutoff between the high- and mid-mobility zones was set as the average behavior of actin-associated proteins, and the low-mobility cutoff of 125 nm corresponds to 2.5 times the average spot localization precision of 50 nm (Supplemental Figure S3). Across the full datasets grouped in this way, high-mobility events have an average apparent D of 6,050 ± 720 nm^2^/sec, mid-mobility events correspond to apparent D of 1,100 ± 14 nm^2^/sec, and low-mobility events correspond to apparent D of 67 ± 35 nm^2^/sec.

As expected, each protein sampled all three behaviors to some extent but membrane-associated proteins predominantly sample the low-mobility class. However, there are some notable exceptions and differences between proteins: Act1p contains a larger population of long-lived, low-mobility events than other actin-binding proteins; Clc1p and End4p contain a population of high-mobility displacements not observed in other membrane-bound proteins Wsp1p, Myo1p, and Pil1p. It is also important to compare the absolute displacements as well as the fractions of events in each class. For example, although Wsp1p has a similar fraction of high-mobility counts as Clc1p, Wsp1p does not contain the population of large-displacement (>500 nm) motions seen in Clc1p and other vesicle-associated proteins and instead the high-mobility events are almost exclusively comprised of short time steps (Figure 6).

## Discussion

### Endocytic proteins display short residence times and rapid turnover

Our single-molecule tracking results indicate that actin and other endocytic proteins turn over multiple times during the overall lifetime of a CME event. This behavior was predicted based on modeling (Berro et al., 2010) and indirect measurements such as FRAP (Kaksonen et al., 2003; Kaksonen et al., 2005), but has never been directly observed in CME before. Other actin structures such as lamellipodia generate sustained force by “treadmilling”, characterized by continuous assembly and disassembly and retrograde flow of incorporated monomers (Pollard and Borisy, 2003; Smith et al., 2013b; Watanabe and Mitchison, 2002). Previous discussions of CME often assumed that the entire actin meshwork assembles and disassembles once during the total lifetime of the patch, with “turnover” referring to the release of the component proteins to be re-used in a later CME event. These models of a single assembly and disassembly of all actin monomers (and actin-associated proteins) would result in an average monomer residence time of ~10 sec (Figure 1). We report actin and associated molecules have residence times most often around 1 to 2 sec, indicating the meshwork instead turns over up to five times, multiplying the total number of molecules that participate, over the lifetime of a single endocytic event.

The characteristic shape of these residence time distributions, a peak with a skewed tail, indicates that the unbinding dynamics of these proteins do not follow a simple kinetic mechanism. Mass action kinetics predict an exponential distribution for single-step unbinding mechanism. In general, multi-step kinetic pathways result in residence times (or “dwell times”) following a gamma distribution (Floyd et al., 2010). Correspondingly, the residence times of actin monomers in a treadmilling filament have been shown to follow complex kinetics resulting in a peak shape that depends on all of the rates of polymerization, aging, cofilin binding, and severing steps (Roland et al., 2008). Although we have not extracted specific rate constants from our data, the shapes of the residence time distributions do appear similar to these models. The skewed peak shape also argues strongly against the interpretation that these single-molecule events arise from transient binding at CME sites or non-productive associations with stable assemblies or other artifacts, processes which would instead result in exponential dwell time distributions according to mass-action kinetics.

We expect that an actin-bound protein would appear to obey similar kinetics as the actin monomer to which it is bound, because the unbinding rates measured for these proteins in vitro are typically far slower than the timescales observed here. For example, unbinding rates from actin filaments have been reported for Acp1p in the range of 10^−4^ sec^−1^ (Bombardier et al., 2015; Kuhn and Pollard, 2007) and for Fim1p in the range of 10^−2^ to 10^−3^ sec^−1^ (Skau et al., 2011). The slight differences between residence times of actin-associated proteins may reflect the delay of binding a filament only after its nucleation (as Fim1p and Acp1p residence times are slightly shorter than Act1p), or the additional time for pre-recruitment before nucleation of the actin branch (as Arc5p residence times are slightly longer than Act1p). The fast sub-population of Fim1p tracks around 0.5 sec likely corresponds to transient binding events that engage one but not both actin-binding domains before unbinding. Such a prominent population of transient events is absent in other proteins. It is interesting that non-actin associated proteins also have residence times with similar shape and timescale, which can be explained by other unique combinations of rates governing those proteins’ assembly and dissociation with the membrane and various binding partners, especially in oligomeric lattice assemblies such as the clathrin coat.

The short lifetimes we measured for clathrin (Clc1p) agree with previous reports that individual clathrin triskelia unbind during CCP development in order to achieve the necessary change in curvature (Avinoam et al., 2015). Previous studies also reported that WASp rapidly exchanges at sites of actin nucleation (Weisswange et al., 2009), although these dynamics have not been measured directly in CME. We showed previously with related methods that a subset of Pil1p molecules undergo turnover at the ends of the linear eisosomes (Lacy et al., 2017), demonstrating the ability of sparse labeling to uncover single-molecule dynamics in dense cellular assemblies.

The rates we report for turnover for the actin meshwork are compatible with previous reports describing the size and speed of actin meshwork polymerization in CME. Kaksonen and colleagues used FRAP to determine that the polymerized actin moves away from the cell membrane at approximately 50 nm/sec (Kaksonen et al., 2003), while single-molecule tracking measurements of actin in lamellipodia report retrograde flow around 30 nm/sec (Yamashiro et al., 2014). Electron microscopy studies have shown that the size of the endocytic actin meshwork extends about 100 to 250 nm from the cell membrane (Idrissi et al., 2012; Kukulski et al., 2012). These measurements, along with the partly overlapping presence of actin assembly and disassembly factors in endocytic patches, have suggested that the meshwork could travel that distance and turn over as rapidly as every 3 to 5 seconds (Goode et al., 2015). Our data are consistent with those estimates, directly showing that actin and associated proteins have typical single-molecule residence times around 1 to 2 seconds, and indeed rarely reside at an endocytic site longer than 3 seconds (Figure 3).

### Motions of endocytic proteins are heterogeneous and dynamic

The apparent diffusion coefficients for actin meshwork proteins and clathrin are similar to previous measurements of vesicle diffusion (2,000 to 8,000 nm^2/sec, (Berro and Pollard, 2014)). It is interesting that in this aspect, the average single-molecule behavior mirrors the bulk behavior of the endocytic patch. However, Wsp1p, Myo1p, and End4p, which have previously been shown to bind at the membrane (Arasada et al., 2018; Mund et al., 2018; Picco et al., 2015; Sirotkin et al., 2010), are still more mobile than Pil1p. This difference may be due to nanoscale motions within the endocytic site, which are a promising target for future study with higher-resolution imaging approaches. The longer distance traveled by long-lived tracks of actin-associated proteins (Figure 4C) can be attributed to molecules that remain associated with the endocytic patch after scission of the vesicle. However, even for those proteins that are known to remain associated with the diffusing vesicle, the large majority of events have very short end-to-end distance (Figure 4A-B). This apparent over-representation of events with restricted motions is consistent with a switch of behavior in the CME assembly from rapid turnover with small motions in the developing CCP before scission to reduced recruitment and sustained disassembly around the vesicle after scission.

Deeper investigation of the single-molecule trajectories reveals heterogeneity in the distributions of displacements over time and unique relationships with track lifetimes (Figure 4 and 6). The high-, mid-, and low-mobility behaviors are consistent with vesicle diffusion (apparent D ~6,000 nm^2^/sec), motions within the CCP-associated meshwork (apparent D ~1,100 nm^2^/sec), and membrane-bound (apparent D ~65 nm^2^/sec) states, respectively. We note that even the highest-mobility events have apparent diffusion coefficient much lower than would be expected for freely diffusing proteins in the cytoplasm (~10 µm^2^/sec, (Milo et al., 2010)), indicating the tracks we observe specifically correspond to proteins incorporated into the large CME assembly and not to free molecules in the background. Moreover, our camera frame rate is not fast enough to capture freely diffusing molecules.

Several key differences are apparent between proteins that display similar MSDs. For example, Act1p and Arc5p sample a long-lifetime, low-mobility behavior not present in Acp1p and Fim1p, perhaps related to pre-nucleation recruitment of Arp2/3 complex and actin monomers by membrane-bound complexes (Mullins et al., 2018; Rohatgi et al., 1999; Smith et al., 2013a). Because these long-lived, immobile actin species have much longer lifetimes than any observed Wsp1p events, they are likely not exclusively bound to Wsp1p but rather to the other membrane-associated nucleation-promoting factors Myo1p (which does display some long-lived, immobile events) or verprolin (Lee et al., 2000; Sirotkin et al., 2005; Sun et al., 2006) or other complexes that remain to be identified. We considered the possibility that these long-lived, low-mobility Act1p steps might correspond to unproductive CCPs or actin filament fragments that remain associated with the base membrane after vesicle scission, but the absence of these events in Acp1p and Fim1p argue against that interpretation. Future study with two-color tracking or other methods would be needed to identify specific complexes within the CME site.

Among membrane-bound proteins, while End4p has a lower MSD than Clc1p, Myo1p, and Wsp1p (Figure 5), both End4p and Clc1p contain some tracks sampling a high-mobility behavior absent in Myo1p and Wsp1p, representing membrane-bound proteins diffusing with the vesicle after scission. This interpretation is consistent with previous reports indicating that the clathrin coat and other membrane-associated proteins including End4p (Sla2 in budding yeast) are internalized with the vesicle, Myo1p remains at the plasma membrane, and Wsp1p moves inward with the CCP tip but mostly dissociates prior to vesicle scission (Arasada et al., 2018; Picco et al., 2018; Picco et al., 2015; Sirotkin et al., 2010). However, while both End4p and Clc1p are known to bind the membrane at endocytic sites long before CCP invagination and scission (bulk lifetimes around 40 sec and 110 sec, respectively (Sirotkin et al., 2010)), Clc1p has considerably fewer long-lived immobile events than End4p, suggesting that it turns over throughout the CCP lifetime and not exclusively during membrane invagination. For all proteins, very long-timescale events (>10 sec) generally exhibit only short displacements, indicating that the molecules are restricted to small motions on the CCP membrane and are disassembled from the diffusing CCV relatively quickly.

### Limitations to current method and data

Single-molecule tracking is a powerful technique to uncover molecular mechanisms in complex systems. We acknowledge that the results of single-molecule tracking are sensitive to the chosen tracking parameters (Jaqaman et al., 2008; Smith et al., 2011), especially because the images’ low signal-to-background ratio results in some missed detections of spots. However, even across a broad range of parameters that introduce verifiable artifacts (see Supplemental Methods), the average single-molecule residence times remain below 3 to 4 sec (Supplemental Figure S4), still much faster than the ~10 sec prediction for single-turnover mechanisms (Figure 1).

### Implications of actin turnover for force production models

Our results have important qualitative and quantitative implications for models of the endocytic machinery, supporting greater total force generation by the actin meshwork than has been previously estimated and enabling network remodeling. If more total molecules of actin are consumed by the endocytic meshwork, the total chemical energy of polymerization through ATP hydrolysis must be several times higher than has been previously estimated. To accumulate the known peak number of actin molecules while sustaining such a high rate of turnover (consuming up to five times more actin and other molecules), the rates of polymerization and disassembly must both be higher than previously estimated. We also observed fast turnover of the actin-nucleation promoting factors Wsp1p and Myo1p. Repeated recruitment for multiple cycles of filament nucleation could support a higher number of polymerizing ends over the lifetime of the CME event, thus multiplying the total load limit of the polymerizing actin meshwork. A recent report proposed that myosin Is (Myo3 and Myo5 in budding yeast) enable higher rates of actin polymerization by pushing the filament ends away from the membrane (Manenschijn et al., 2018). Another recent report demonstrated that the mechanical stress in a highly-crosslinked actin network can greatly enhance the rate of ADF/cofilin severing of actin filaments (Wioland et al., 2019), which could also support the high turnover rates we observe in CME patches. Previous quantitative models should be reconsidered with these changes to the assumed rates, and novel experimental methods may be needed to measure rates and molecular complexes at the endocytic site.

Fast turnover of the actin meshwork components could affect other higher-order mechanisms of force generation. For example, continuous unbinding of fimbrin crosslinkers would allow remodeling of the meshwork and spatiotemporal redistribution of energy stored in crosslinkers (Ma and Berro, 2018; Picco et al., 2018). In addition to allowing energy to be stored and released, such remodeling of the actin meshwork could enable important changes in the orientation of filaments and their force outputs over time. Further experimental and modeling work will be needed to clarify how sustained turnover affects the various mechanochemical mechanisms of force generation in CME.

Future studies of CME should account for the idea that the endocytic actin meshwork and membrane coat are highly dynamic, turning over multiple times to generate, distribute, and transmit the forces that deform the plasma membrane into a vesicle. We expect that the increasing accessibility of super-resolution microscopy and single-molecule techniques as well as added levels of detail in computational models will enable further exploration of the molecular mechanisms of CME and other complex molecular assemblies in live cells.

## Acknowledgements

We thank Ronan Fernandez for assistance in creating yeast strains, Olivier Trottier for assistance with an earlier version of the Matlab analysis scripts, and members of the Berro lab for helpful discussions. This research was supported in part by National Institutes of Health/National Institute of General Medical Sciences Grant R01GM115636. MML was supported by National Institutes of Health Training Grant T32GM008283. We also acknowledge support from the Raymond and Beverly Sackler Institute for Biological, Physical and Engineering Sciences at Yale University.

## Author Contributions

Conceptualization was by MML and JB; Methodology by MML, DB, and JB; Investigation by MML; Software by MML and DB; Formal Analysis by MML; Writing – Original Draft by MML; Writing – Review & Editing by MML, DB, and JB; Visualization by MML; Resources by DB and JB; Funding Acquisition by JB; and Supervision by DB and JB.

### Declaration of Interests

The authors declare no competing financial interests. MML is a current employee of Elsevier.

## Materials and Methods

### Model simulations

Simulated profiles of numbers of molecules and binding/unbinding events were calculated with Matlab (Mathworks), as lists of timestamps generated according to each type of dynamic model. The simulation script is given as Supplemental File 1.

For “Random assembly” models, binding times were randomly chosen according to a normal distribution centered at one quarter of the total patch lifetime and standard deviation of one tenth of the total lifetime. For “Random assembly/Random disassembly”, the unbinding times were randomly chosen according to a normal distribution centered at three quarters of the total lifetime and standard deviation of one tenth of the total lifetime. For “Random assembly with First-in/First-out disassembly”, binding and unbinding times were chosen on the same distributions as above, but were sorted so that the corresponding values of binding times and unbinding times could be matched in order from lowest-to-highest. Single monomer residence times were calculated by subtracting each unbinding time from its corresponding binding time.

For “Continuous turnover” model, binding times were randomly chosen according to a normal distribution, taking the absolute value of a distribution centered at one third of the total lifetime and standard deviation of one fifth of the total lifetime. Single monomer residence times were chosen according to a gamma distribution with shape parameter equal to 2 and scale parameter equal to 1, to represent a multi-step dwell time kinetic giving a shape and mean resembling the observed lifetime distributions. Unbinding times were calculated by taking the sum of the binding time and a randomly-chosen residence time.

The profile of number of molecules present in the patch at each 1-sec time bin was calculated as the number of molecules with binding time smaller than or equal to the current time bin and unbinding time larger than the current time bin. Simulations were performed with 900 total molecules in a lifetime of 20 sec, repeated 50 times for each model.

### Yeast strains

We generated *Schizosaccharomyces pombe* strains with SNAP-tag (Addgene plasmid 87024) or mEGFP (Addgene plasmid 87023) inserted at the genomic loci of various proteins, using either homologous recombination with kanamycin selection (Bahler et al., 1998) or CRISPR-Cas9 gene editing with fluoride selection (Fernandez and Berro, 2016). Because *S. pombe* is inviable when its sole source of actin is fused with a fluorescent protein (Wu et al., 2008; Wu and Pollard, 2005), we integrated SNAP-act1 into the *leu1*+ locus under control of the *41nmt* promoter, as previous studies have done for mEGFP-act1. We did not measure the expression level of SNAP-Act1p or its effect on actin functionality, but we expect it to behave similarly to mEGFP-Act1p used previously. Previous studies using this strategy report the actin fusion protein represents around 5% of the total actin in the cell (Berro and Pollard, 2014; Sirotkin et al., 2010; Wu et al., 2008; Wu and Pollard, 2005). The strains used in this study are listed in Supplemental Table S1.

### Growth and SNAP-tag labeling

Cells were grown in liquid YE5S medium at 32° C to exponential phase (OD_595_ between 0.4 and 0.6) then diluted into liquid EMM5S medium and grown for 12 to 24 hours at 25° C before labeling with SNAP fluorophore (Keppler et al., 2003; Lukinavicius et al., 2015; Stagge et al., 2013). As discussed previously (Lacy et al., 2017), the SNAP-substrate fluorophore does not accumulate in high amounts in yeast cells containing multidrug exporter genes and intact cell walls. Cells were diluted to 0.1 OD_595_ in 1 mL of EMM5S containing 1 µM of silicon-rhodamine benzylguanine derivative SNAP-SiR647 (SNAP-Cell^®^ 647-SiR, New England Biolabs). Culture tubes were wrapped in aluminum foil to protect them from light and incubated on a rotator overnight (about 15 hours) at 25° C. Cells were washed five times by centrifuging at 1,500xg for 3 minutes and resuspending in 1 mL of EMM5S, then incubated at 25° C for an additional hour, then washed five times again by centrifuging at 1,500xg for 3 minutes and resuspending in 1 mL of EMM5S. Cells were finally resuspended in 20 to 100 µL of 0.22-µm filtered EMM5S to achieve suitable density for imaging. SNAP-tag labeling efficiency was not determined before tracking analysis, but is estimated to be around 0.1% to 1% of the SNAP-tag fusion protein’s expression level.

Cells expressing mEGFP fusion were grown in YE5S and EMM5S liquid media as above, then washed once by centrifuging at 1,500xg for 3 minutes and resuspending in 20 to 100 µL of 0.22-µm filtered EMM5S to achieve suitable density for imaging.

### Paraformaldehyde fixation

For imaging fixed cells, SNAP-tag expressing cells were grown and labeled and washed as above. Cells were incubated in 3.6% paraformaldehyde solution for 15 minutes on a rotator at room temperature. Cells were then pelleted by centrifugation at 10,000xg for 1 minute and washed three times by resuspending in filtered EMM5S and centrifuging at 1,500xg for 3 minutes. Cells were finally resuspended in 20 to 100 µL of 0.22-µm filtered EMM5S to achieve suitable density for imaging.

### Imaging

Cells were pipetted on to pads of 0.25% gelatin prepared with 0.22-µm filtered EMM5S, covered with a #1.5 coverslip, which had been washed in ethanol for 30 minutes and plasma cleaned for 3 minutes to avoid nonspecific dye or other autofluorescent particles on the surface, and sealed around the edges with Valap. Samples were imaged on an Eclipse Ti inverted microscope (Nikon) equipped for through-objective TIRF, with a 642 nm excitation laser for SiR imaging (Spectral Applied Research) and a 488 nm laser for mEGFP imaging (Spectra-Physics). Images were recorded through a 60x/1.49 NA Apo TIRF objective (Nikon) and further magnified with the microscope’s 1.5x lens, and detected using an iXon DU897 EMCCD camera (Andor); image pixels correspond to 178 nm. The microscope, camera, and illumination were controlled through Nikon Elements software.

For single-molecule tracking in SNAP-SiR labeled samples, the imaging focal plane is set about 1 to 1.5 μm below the cell midplane, just above the base of the cells adjacent to the coverslip, and the laser is angled for near-TIRF illumination, so that spots at the membrane are in focus and cytoplasmic fluorescence is reduced. The 642 nm laser illumination intensity was 0.5 or 0.8 W/cm^2^ (measured exiting the objective). The camera was set to 100 msec exposure, with EM gain set to 300 at 5 MHz readout with 14-bit digitization depth. We recorded 60-second movies, starting recording with the laser off and turning it on after a few frames so that the initial fluorescence signal is not lost due to hardware delay times.

For endocytic patch tracking in samples expressing mEGFP, the 488 nm laser illumination was 0.2 W/cm^2^, with camera exposure 100 msec and EM gain 300. We recorded 120-sec movies, starting with the laser off and turning it on after a few frames.

For two color imaging, we collected alternating red-channel and green-channel images, switching between 642 nm laser (0.8 W/cm^2^) and 488 nm laser (0.2 W/cm^2^) with 200 msec exposure time and EM gain set to 300, with about 1 sec delay between acquisition frames to switch filter sets. A sample of Tetraspeck beads (0.2 μm) was prepared and imaged using the same protocol to ensure alignment between channels.

### Image analysis – single molecule localization and tracking

Super-resolution spot localization and track generation were performed in the Python Microscopy Environment (PYME) (http://www.python-microscopy.org, (Baddeley et al., 2011)). Metadata for the images were determined as follows: camera read noise (88.8), from manufacturer’s specifications; camera noise factor (1.41), standard correction for EMCCD cameras; electrons per count (49) and true EM gain (167), by calibration of blank movies recorded at varying intensity and gain settings, following (Hirsch et al., 2013); camera AD Offset (105), by measuring the average pixel intensity of dark frames before activating the laser. Each movie was processed for spot detection using a 2D Gaussian model with local intensity scaling threshold factor of 0.8; “debounce rad”, or the distance (in pixels) within which two fluorophores cannot be distinguished as 3; and the temporal background subtraction feature turned off. The local background and total spot intensity are both fitted with 2D Gaussians and subtracted to determine the spot intensity.

Candidate spots were filtered based on spot intensity (20 to 200 AU), spot size sigma (100 to 300 nm), and localization precision in x and y (0 to 150 nm). Single-molecule trajectories were generated by linking spots in successive frames within a limit of 100 nm, and allowing a maximum gap of 6 frames (spanning missed localizations in up to 4 frames), as discussed in Supplemental Methods. Any spots which are not incorporated into a track of at least three spots, and any tracks which appear outside the cells, are discarded. Localization and tracking results were exported as text files to be further analyzed in Matlab.

Spots in the fixed sample of Acp1p-SiR were analyzed using TrackMate plugin (Tinevez et al., 2017) for FIJI (Schindelin et al., 2012; Schneider et al., 2012), using spot detection with the Difference of Gaussians detector, threshold 2.5, maximum diameter 1 µm, and median filtering, and tracked using semi-automated manual tracking. We did not apply strict limits to spot quality and signal thresholds but rather relied on visual inspection and automated detection of spots’ signal in successive frames, allowing gaps of missed localizations up to 4 frames.

### Image analysis – GFP patch tracking

We tracked GFP spots with the TrackMate plugin (Tinevez et al., 2017) for FIJI (Schindelin et al., 2012; Schneider et al., 2012), using spot detection with the Difference of Gaussians detector, threshold 2.0, maximum diameter 1 µm, and median filtering, and tracked using semi-automated manual tracking. We visually confirmed that tracks did not overlap with neighboring spots and spanned the full lifetime of intensity increase and decay, rather than a sharp disappearance due to diffusing out of the focal plane. Only these manually-curated tracks were exported for further analysis in Matlab.

Spots in the two-color images were identified and scored manually. Spot detection and colocalization algorithms would have difficulty because the GFP patches vary widely in intensity.

### Data analysis

Tracking data were read, curated, and analyzed with custom-written code in Matlab (Mathworks). We excluded tracks with lifetime shorter than 0.3 seconds and tracks which began before the start-time cutoff determined for each protein (see Supplemental Figure S5). Because the first few frames of the movies are more crowded (potentially introducing errors in track linking) and may contain molecules already present in CME structures (yielding only partial information on the molecule’s true trajectory) or other artifacts (e.g. high background or molecular aggregates that have not been photobleached), tracks which started within the cutoff time after the laser was turned on were discarded. We chose the appropriate start-time cutoffs for each protein dataset as the first time point where the difference between the current lifetimes distribution and the distribution using the next cutoff (1 sec later) was not significant by KS test.

A variety of features are calculated from the timestamp and position data such as the lifetime, stepwise displacements, velocity, and other derivative characteristics at the level of single spots or tracks. Statistics for numbers of tracks, movies analyzed, and independent samples prepared are given in Table 1. Single-molecule tracking datasets for endocytic proteins typically represent 10 to 30 movies analyzed from between two to five independent samples. Standard deviation of single-molecule residence time data was determined by calculating the mean track lifetimes from each movie individually, and computing the standard deviation of these separate results. Where indicated, we used a two-sample Kolmogorv-Smirnov test for statistical significance of differences between distributions.

### Software and reagents availability

The Python microscopy environment (PYME) software is available at www.python-microscopy.org, and our Matlab analysis script is available as Supplemental Files. We will happily share any S. pombe strains upon request and plasmids are available through Addgene.

